# Enhanced replication of contemporary human highly pathogenic avian influenza H5N1 virus isolate in human lung organoids compared to bovine isolate

**DOI:** 10.1101/2024.08.02.606417

**Authors:** Meaghan Flagg, Brandi N. Williamson, Johan A. Ortiz-Morales, Tessa R. Lutterman, Emmie de Wit

**Affiliations:** National Institute of Allergy and Infectious Diseases, National Institutes of Health, Hamilton, MT, USA

**Keywords:** influenza A virus, highly pathogenic avian influenza, H5N1, clade 2.3.4.4b, outbreak, human lung organoids

## Abstract

We compared virus replication and host responses in human alveolar epithelium infected with highly pathogenic avian influenza (HPAI) H5N1 viruses. A/Vietnam/1203/2004 replicated most efficiently, followed by A/Texas/37/2024, then A/bovine/Ohio/B24OSU-342/2024. Induction of interferon-stimulated genes was lower with A/Texas/37/2024 and A/bovine/Ohio/B24OSU-342/2024, which may indicate a reduced disease severity of these viruses.

---

Clade 2.3.4.4b highly pathogenic avian influenza (HPAI) H5N1 viruses have circulated in avian species in North America since 2022. Subsequently, these viruses have been detected in a wide range of mammalian species (1). In 2024, clade 2.3.4.4b HPAI H5N1 virus was detected in dairy cattle, in both tissue samples and milk from infected animals (2), and has subsequently spread to multiple herds around the United States. The broadened host range of clade 2.3.4.4b H5N1 viruses along with unprecedented levels of transmission between mammals has raised concerns about potential spillover into humans.

As of July 25, 2024, 13 human cases of HPAI H5N1 virus infection have been confirmed in the United States (3). Several of these cases are linked to exposure to infected cattle. However, recent outbreaks in Colorado have resulted in identification of additional human cases linked to infected poultry (3). Virus isolated from a worker at a Texas dairy farm (A/Texas/37/2024) was shown to be closely related to viruses circulating in cattle, and it is presumed that this case is a result of direct cow-to-human transmission (4). Reported symptoms included conjunctivitis, as well as mild respiratory symptoms in one case (5).

This is in stark contrast to prior cases of HPAI H5N1 virus infection in humans, which resulted in severe respiratory disease and mortality rates upwards of 50% (6). In order to assess the risk of developing severe disease following infection with contemporary HPAI H5N1 virus, we evaluated virus replication, host cell survival, and induction of innate immune responses in human alveolar epithelium infected with A/Texas/37/2024 or cattle isolate A/bovine/Ohio/B24OSU-342/2024, compared to a historical H5N1 isolate A/Vietnam/1203/2004, which was derived from a fatal human case (7).

## The Study

Virus replication and host cell damage in the alveolar epithelium are key drivers of severe respiratory disease. Human lung organoids are a physiologically relevant state-of-the-art model of primary human alveolar epithelium. Lung organoids consisting of alveolar type 2 (AT2) epithelial cells can be cultured from adult stem cells isolated from lung tissue (8,9), or from induced pluripotent stem cells (iPSCs) differentiated into AT2 cells (10). Both model systems accurately recapitulate the fitness and pathogenicity of respiratory viruses as observed in humans (8,9,11,12). We infected AT2 cells from both iPSC-derived and adult stem cell-derived human lung organoids (ihLOs and hLOs, respectively; human donor lung tissue was kindly provided by Chuong D. Hoang and Nathanael Pruett at the National Cancer Institute. De-identified human lung tissue samples were collected in accordance with Institutional Review Board-approved protocols at the NIH Clinical Center) with three HPAI H5N1 isolates and compared virus replication, host cell survival, and innate immune responses over time.

In both ihLOs and hLOs, influenza virus A/Vietnam/1203/2004 replicated faster and to higher titers (Figure 1). Replication of the cattle isolate A/bovine/Ohio/B24OSU-342/2024 (kindly provided by Richard Webby, St. Jude’s Children’s Research Hospital, and Andrew Bowman, Ohio State University) was significantly lower compared to the A/Vietnam/1203/2004 isolate. Notably, we detected a trend towards increased replication of A/Texas/37/2024 (kindly provided by Todd Davis, Centers for Disease Control and Prevention), compared to the bovine isolate, suggesting enhanced fitness of this virus in human cells compared to its predecessors circulating in cattle.

**Figure 1.**
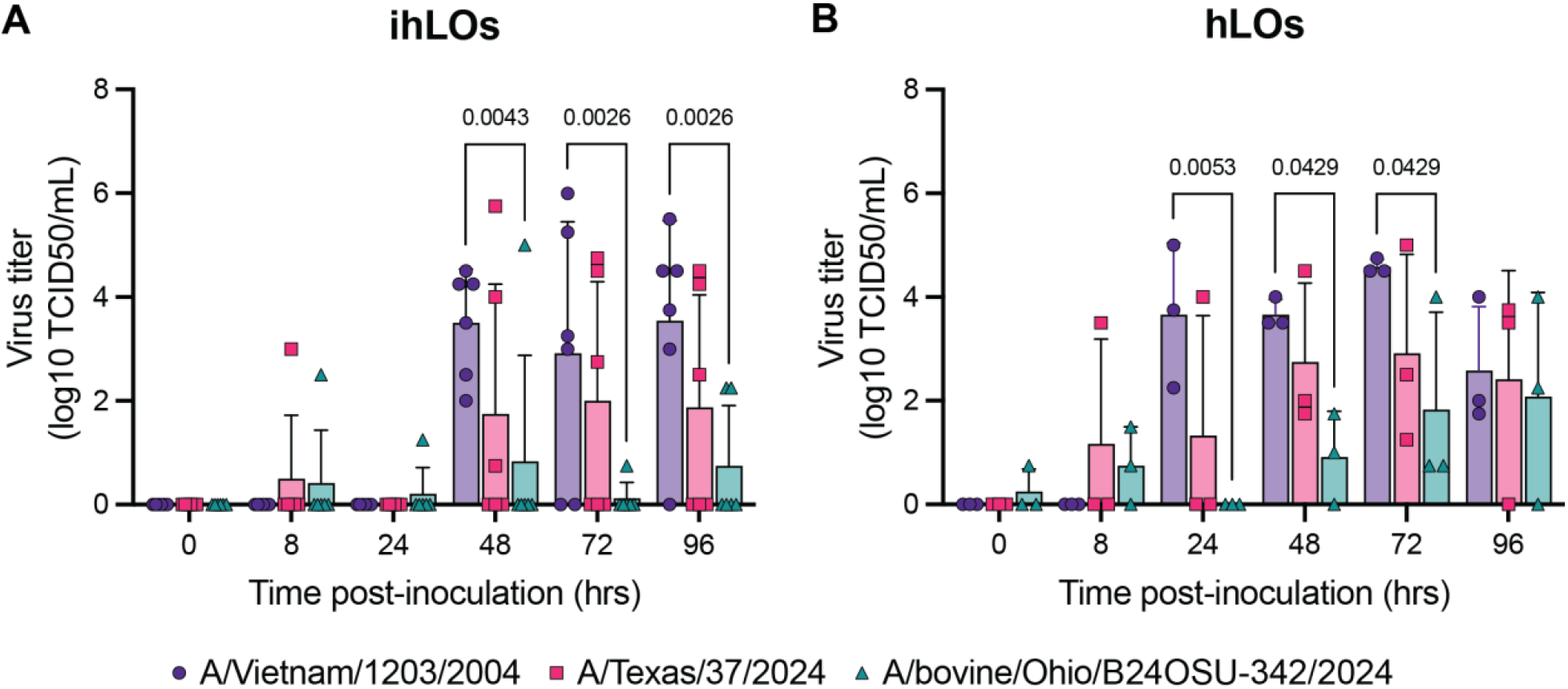
A/Vietnam/1203/2004 virus replicates most efficiently in human lung organoids. A) iPSC-derived human lung organoids (ihLO) or B) adult-derived human lung organoids (hLO) were infected at an MOI of 0.1 with one of the three indicated HPAI H5N1 influenza A virus isolates. Samples were taken at 0, 8, 24, 48, 72, and 96 hrs post-inoculation and titered on MDCK cells. Titrations were read after 3 days using a hemagglutination assay using turkey red blood cells. Error bars denote mean and standard deviation. Statistical analysis was conducted using two-way ANOVA followed by Tukey’s post-test. P-values <0.05 are indicated.

To evaluate potential pathogenicity of the contemporary HPAI H5N1 viruses, we quantified cell death over time in infected lung organoids. Cell death was observed earlier in organoids infected with the A/Vietnam/1203/2004 isolate (Figure 2). Infection with the A/Texas/37/2024 and A/bovine/Ohio/B24OSU-342/2024 isolates also resulted in cell death, although at later timepoints (72-96hr post-inoculation). This is in accordance with the slower replication kinetics observed with these viruses compared to the A/Vietnam/1203/2004 isolate (Figure 1).

**Figure 2.**
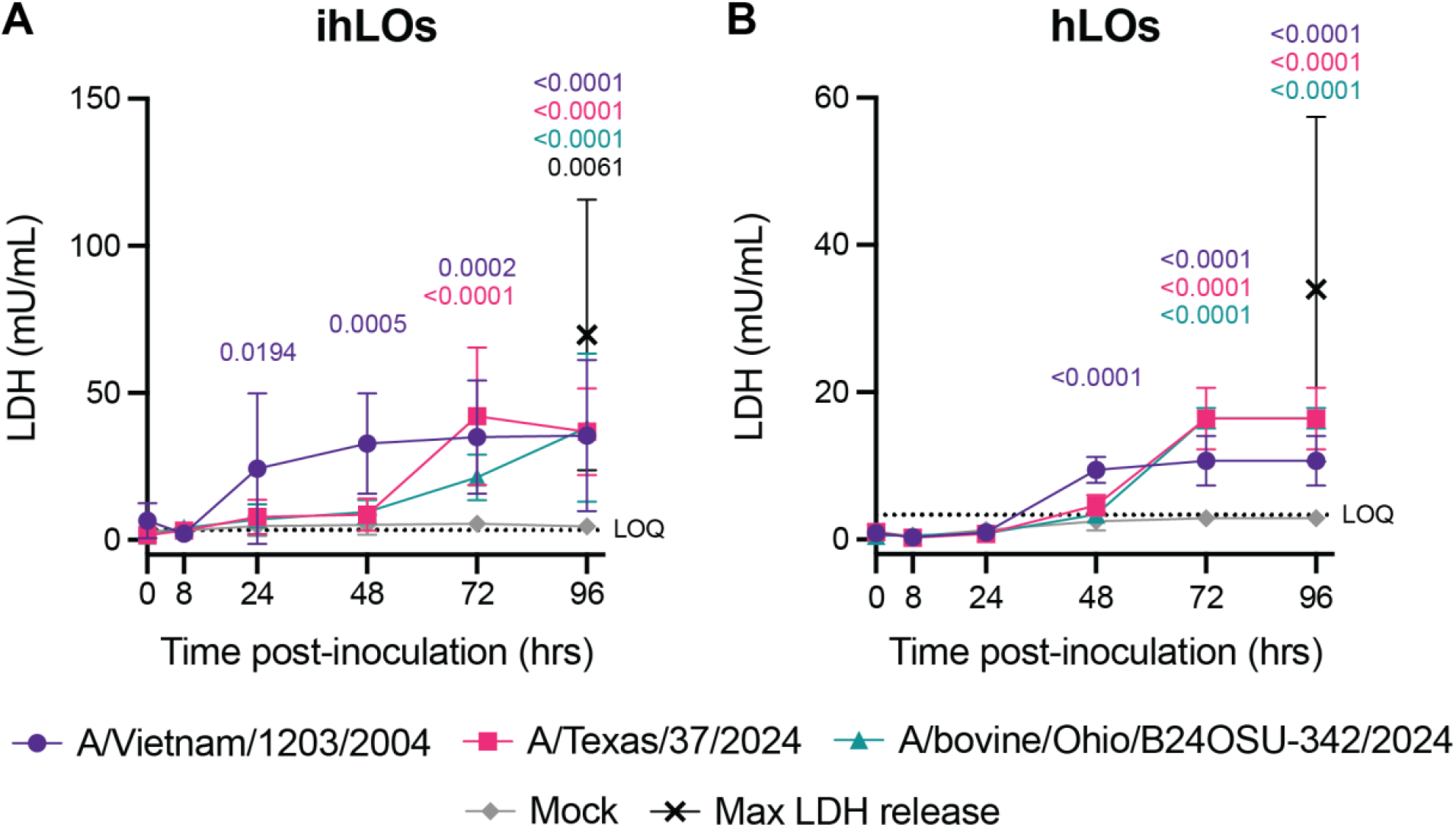
Cytotoxicity of HPAI H5N1 virus infection in human lung organoids. A) iPSC-derived human lung organoids (ihLO) or B) adult-derived human lung organoids (hLO) were infected as described in Figure 1. Samples taken at 0, 8, 24, 48, 72, and 96hrs post-inoculation were collected and tested for release of lactate dehydrogenase into culture supernatant according to the LDH-Glo assay protocol (Promega) as an indicator of cell death. Results were converted to mU/mL LDH determined by the standard curve using a simple linear regression. Dashed line indicates lower limit of quantification. Cells lysed with Triton X-100 were included as maximum LDH release controls. Error bars denote mean and standard deviation. Statistical analysis was conducted using two-way ANOVA followed by Dunnett’s post-test. P-values indicate comparison to mock-infected controls; p-values less than 0.05 are shown.

We quantified induction of interferon-stimulated genes (ISGs) IFITM3, ISG15, and ISG20 by qRT-PCR as a measure of the host innate immune response to infection (Figure 3). ISG induction was highest in organoids infected with the A/Vietnam/1203/2004 isolate. This was most pronounced in the adult stem cell-derived hLOs, where ISG induction was not detected in organoids infected with the A/Texas/37/2024 or A/bovine/Ohio/B24OSU-342/2024 isolates, despite the presence of replicating virus. Moderate induction of IFITM3 and ISG15 was detected in ihLOs infected with the A/Texas/37/2024 and A/bovine/Ohio/B24OSU-342/2024 isolates.

**Figure 3.**
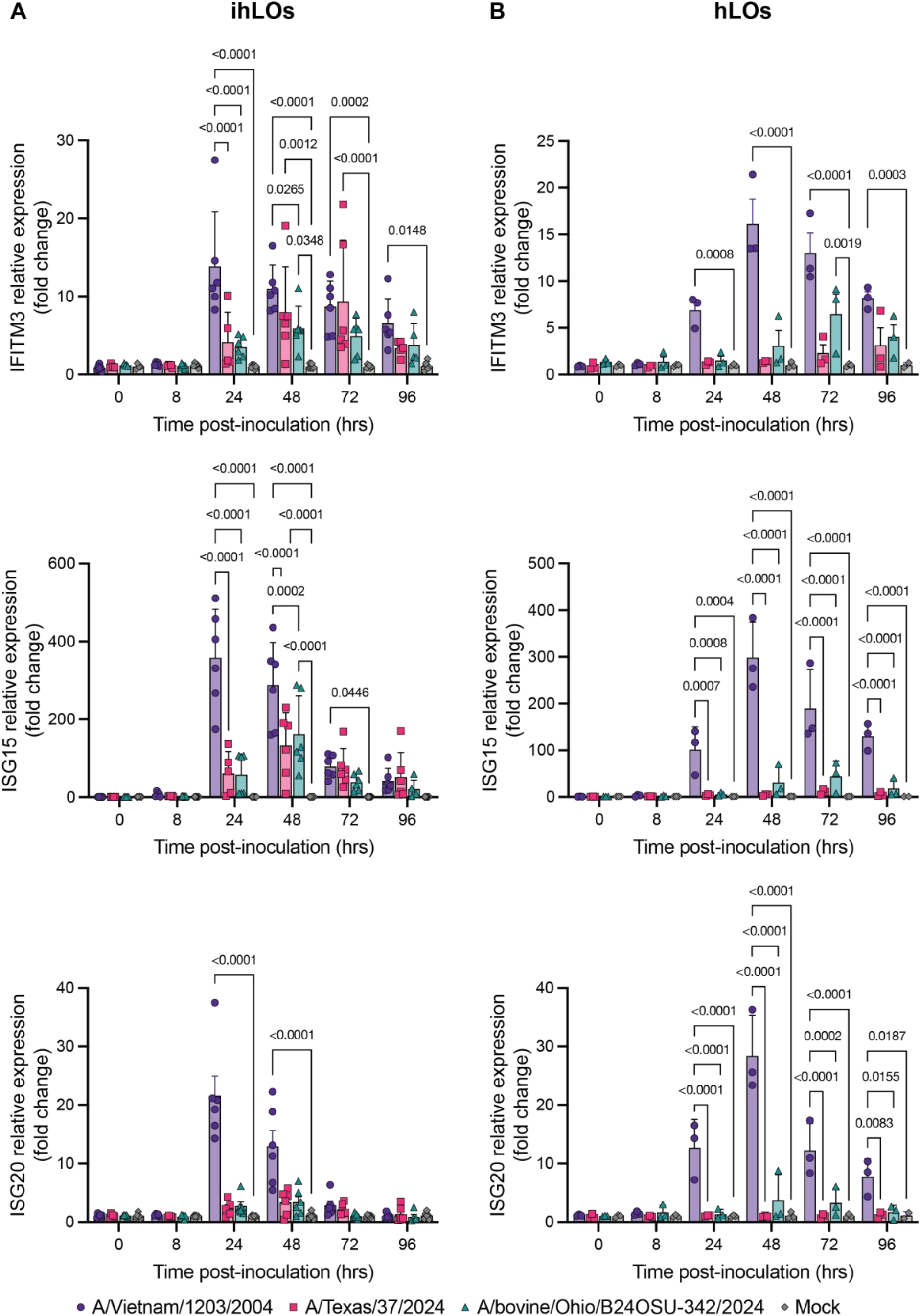
Induction of interferon stimulated genes (ISG) in HPAI H5N1 virus-infected human lung organoids. (A-C) iPSC-derived human lung organoids (ihLO) or (D-F) adult-derived human lung organoids (hLO) were infected as described in Figure 1. RNA from 2.5×10^4^ cells was extracted using the QIAGEN RNeasy kit (QIAGEN) following the tissue extraction instructions. Quantitative reverse transcription PCR (qRT-PCR) was run using commercially available primers (IDT) to ISG15, ISG20, and IFITM3. Data were normalized to timepoint-matched mock-infected controls and fold induction is shown. Error bars denote mean and standard deviation. Statistical analysis was performed using two-way ANOVA followed by Dunnett’s post-test; p values less than 0.05 are shown.

## Conclusions

The unusual transmission of clade 2.3.4.4b HPAI H5N1 viruses between mammals has raised concerns about the risk of spillover into the human population, and the possibility of outbreaks leading to severe disease. We assessed virus replication and host responses in human alveolar epithelium, since virus replication and host cell damage in this site is a key driver of severe respiratory disease. The reduced replication levels of the A/Texas/37/2024 and A/bovine/Ohio/B24OSU-342/2024 isolates in lung organoids compared to the A/Vietnam/1203/2004 isolate may explain why recent human cases involving the clade 2.3.4.4b viruses resulted in mild illness (4,6), as opposed to the severe respiratory disease associated with previous outbreaks in Vietnam (13,14). The presence of a lysine at position 627 in the PB2 protein has been associated with adaption of avian influenza viruses to mammalian hosts and is known to increase virus replication in the mammalian respiratory tract (15). This substitution (E627K) is present in both the A/Vietnam/1203/2004 and A/Texas/37/2024 viruses but not the A/bovine/Ohio/B24OSU-342/2024 isolate (4), which may explain the increased replication observed for the A/Texas/37/2024 isolate compared to the bovine isolate.

Another factor contributing to the reduced disease severity in humans following infection with clade 2.3.4.4b viruses compared to previous HPAI H5N1 virus cases may be differential activation of the immune system. We observed significantly higher induction of ISGs in lung organoids infected with the A/Vietnam/1203/2004 isolate. An overly exuberant immune response, including cytokine storm, is known to play a role in the high mortality from HPAI H5N1 virus infections observed during the 2003 and 2004 outbreaks in Vietnam (14). Limited innate immune activation elicited by the A/Texas/37/2024 and A/bovine/Ohio/B24OSU-342/2024 isolates may contribute to their reduced pathogenicity.

Despite differences in virus replication and ISG induction, we observed similar levels of cell death by 96 hours post-inoculation for all three viruses. Previous work has shown that direct virus-induced cytotoxicity is not always indicative of pathogenicity *in vivo*, as cytotoxicity was not observed in SARS-CoV-2-infected lung organoids (11), despite the ability of this virus to cause severe respiratory disease. Taken together, these data suggest that epithelial-extrinsic factors, possibly related to immune activation, govern pathogenicity *in vivo*.

In summary, this study provides a characterization of virus replication and host responses to infection in human alveolar epithelium between a contemporary clade 2.3.4.4b human HPAI H5N1 isolate compared to the highly virulent A/Vietnam/1203/2004 virus. Further studies are warranted to understand how these viruses interact with the innate immune system, and how this affects pathogenesis *in vivo*.

## Acknowledgements

This work was supported by the Intramural Research Program of the National Institute of Allergy and Infectious Diseases, NIH.

## Data availability statement

Data included in this manuscript have been deposited in Figshare at https://doi.org/10.6084/m9.figshare.26487574

## References

1. Elsmo EJ, Wünschmann A, Beckmen KB, Broughton-Neiswanger LE, Buckles EL, Ellis J, et al. Highly Pathogenic Avian Influenza A(H5N1) Virus Clade 2.3.4.4b Infections in Wild Terrestrial Mammals, United States, 2022 - Volume 29, Number 12—December 2023 - Emerging Infectious Diseases journal - CDC. Emerg Infect Dis. 2023;29(12):2451–60.

2. Burrough ER, Magstadt DR, Petersen B, Timmermans SJ, Gauger PC, Zhang J, et al. Highly Pathogenic Avian Influenza A(H5N1) Clade 2.3.4.4b Virus Infection in Domestic Dairy Cattle and Cats, United States, 2024. Emerg Infect Dis. 2024;30(7):1335–43.

3. CDC Newsroom. CDC Confirms Three Human Cases of H5 Bird Flu Among Colorado Poultry Workers [Internet]. 2024. Available from: https://www.cdc.gov/media/releases/2024/s0725-three-human-cases-of-h5-bird-flu.html

4. Uyeki TM, Milton S, Hamid CA, Webb CR, Presley SM, Shetty V, et al. Highly Pathogenic Avian Influenza A(H5N1) Virus Infection in a Dairy Farm Worker. N Engl J Med. 2024;390(21):2028–9.

5. CDC Newsroom. CDC Confirms Second Human H5 Bird Flu Case in Michigan; Third Case Tied to Dairy Outbreak [Internet]. 2024. Available from: https://www.cdc.gov/media/releases/2024/p0530-h5-human-case-michigan.html

6. CDC. Technical Report: June 2024 Highly Pathogenic Avian Influenza A(H5N1) Viruses [Internet]. 2024. Available from: https://www.cdc.gov/bird-flu/php/technical-report/h5n1-06052024.html

7. Maines TR, Lu XH, Erb SM, Edwards L, Guarner J, Greer PW, et al. Avian Influenza (H5N1) Viruses Isolated from Humans in Asia in 2004 Exhibit Increased Virulence in Mammals. J Virol. 2005;79(18):11788–800.

8. Katsura H, Sontake V, Tata A, Kobayashi Y, Edwards CE, Heaton BE, et al. Human Lung Stem Cell-Based Alveolospheres Provide Insights into SARS-CoV-2-Mediated Interferon Responses and Pneumocyte Dysfunction. Cell Stem Cell. 2020;27:890-904.e8.

9. Youk J, Kim T, Evans KV, Jeong YI, Hur Y, Hong SP, et al. Three-dimensional human alveolar stem cell culture models reveal infection response to SARS-CoV-2. Cell Stem Cell. 2020;27(6):905-919.e10.

10. Jacob A, Vedaie M, Roberts DA, Thomas DC, Villacorta-Martin C, Alysandratos KD, et al. Derivation of self-renewing lung alveolar epithelial type II cells from human pluripotent stem cells. Nat Protoc. 2019;14(12):3303–32.

11. Flagg M, Goldin K, Pérez-Pérez L, Singh M, Williamson BN, Pruett N, et al. Low level of tonic interferon signalling is associated with enhanced susceptibility to SARS-CoV-2 variants of concern in human lung organoids. Emerg Microbes Infect [Internet]. 2023;12(2):2276338. Available from: https://doi.org/10.1080/22221751.2023.2276338

12. Dobrindt K, Hoagland DA, Seah C, Kassim B, O’Shea CP, Murphy A, et al. Common Genetic Variation in Humans Impacts In Vitro Susceptibility to SARS-CoV-2 Infection. Stem Cell Rep. 2021;16(3):505–18.

13. Tran TH, Nguyen TL, Nguyen TD, Luong TS, Pham PM, Nguyen van VC, et al. Avian Influenza A (H5N1) in 10 Patients in Vietnam. N Engl J Med. 2004;350(12):1179–88.

14. Peiris J, Yu W, Leung C, Cheung C, Ng W, Nicholls J, et al. Re-emergence of fatal human influenza A subtype H5N1 disease. Lancet. 2004;363(9409):617–9.

15. Hatta M, Hatta Y, Kim JH, Watanabe S, Shinya K, Nguyen T, et al. Growth of H5N1 Influenza A Viruses in the Upper Respiratory Tracts of Mice. PLoS Pathog. 2007;3(10):e133.

